# Dynamic Properties of Simulated Brain Network Models and Empirical Resting State Data

**DOI:** 10.1101/344473

**Authors:** Amrit Kashyap, Shella Keilholz

## Abstract

Brain Network Models have become a promising theoretical framework in simulating signals that are representative of whole brain activity such as resting state fMRI. However, it has been difficult to compare the complex brain activity between simulated and empirical data. Previous studies have used simple metrics that surmise coordination between regions such as functional connectivity, and we extend on this by using various different dynamical analysis tools that are currently used to understand resting state fMRI. We show that certain properties correspond to the structural connectivity input that is shared between the models, and certain dynamic properties relate more to the mathematical description of the Brain Network Model. We conclude that the dynamic properties that gauge more temporal structure rather than spatial coordination in the rs-fMRI signal seem to provide the largest contrasts between different BNMs and the unknown empirical dynamical system. Our results will be useful in constraining and developing more realistic simulations of whole brain activity.

## Introduction

Whole brain activity is complex but critical for understanding behavior and function of the central nervous system. To explain its complexity, studies have used resting state fMRI (rs-fMRI) scans and functional connectivity (FC) analysis to describe the coordination between different brain regions of interest (ROIs) during rest, task, and other behavioral paradigms (Smith 2009). In recent years, analysis of FC data has moved beyond looking at average statistical relationships maintained over the course of a long scan (average FC) to dynamic analysis methods that assume the coordination of brain activity changes on a moment to moment basis (Shakil et al., 2016). The anatomical connections of the brain are assumed to remain constant on these short times scales so that the time-varying coordinated activity plays out over the same framework of structural connectivity (SC) based on white matter connections over time (Cabral et al., 2017; Deco et al., 2017; Shen et al., 2015). Thus, the brain’s activity can be modeled as interactions of ROIs connected by a structural network, where the activity of each ROI is a function of the local state of processing plus a delayed function of the activity of its network neighbors (Breakspear, 2017; Sanz Leon et al., 2015). The resulting set of differential equations form a *dynamical system* which can be used as a generative model to simulate activity across the whole brain for a given state vector of ROI activity. Thus, *dynamic* FC can be thought of as the brain’s trajectory across the phase space of the underlying *dynamical system* (Deco et al., 2013; Cabral et al., 2017).

Numerical simulations of this network of ROIs, known as the brain network model (BNM) (Sanz Leon et al., 2015), simulate spontaneous neural activity in the absence of external stimuli. Without external stimuli, as in all of rs-fMRI, there exists no time-locked measure or event that would allow for straightforward comparison across modalities. Instead, researchers have used measures that summarize activity throughout the brain, such as average FC, estimated through granger causality or correlation and used distance metrics or graph theoretic analysis to quantify how similar they are across modalities (Cabral et al., 2011; Li et al., 2015; Senden et al., 2017). A wide range of BNMs have successfully reproduced the most prominent features of average FC (Cabral et al., 2011; Cabral et al., 2012; Hansen et al., 2014; Senden et al., 2017; Sanz Leon et al., 2015). Newer studies have tried to develop more complex BNMs to describe transient features observed in resting state, such has the switching between two FC states naturally during rest (Hansen et al., 2014; Cabral et al., 2017; Deco et al., 2018). However, as BNMs have become more sophisticated, dynamic analysis methods for rs-fMRI have also become more developed, leading to the question of what dynamical properties of rs-fMRI BNMs can reproduce. Many dynamic analysis methods have been applied to rs-fMRI and provide complementary views of the brain activity (Hutchison et al., 2013; Keilholz et al., 2017). The replication of these dynamic features using a generative model would provide new insight on how they might arise and what they could represent. At the same time, more stringent constraints based on dynamic rather than average rs-fMRI features would provide better discrimination between different models and between different parameterization of the same model.

The following study compares the dynamics observed in rs-fMRI to the results of the same analysis methods applied to two BNMs. We simulate two different types of BNMs with delayed inputs, the Kuramoto oscillator model and the Fokker-Planck model (also called General Linear Model in literature), and then apply four of the most common dynamic analysis techniques to compare features found in the simulated data with those found in rs-fMRI scans (Cabral et al., 2011, 2012). We chose the Kuramoto and the Fokker-Planck model because they were shown to be robust models in reproducing brain activity, have relatively small number of parameters to optimize, and have also shown to exhibit different dynamical properties so we expected see a contrast between them (Deco et al., 2010; Cabral et al., 2017). We chose specific analysis techniques that examine the dynamical properties by testing the signal for states, repeating events or trajectories that are representative of its higher order spatial-temporal structure:

1. **Point-process or neural avalanche theory**, which models the BOLD signal as a combination of discrete neural events or avalanches (Caberello et al., 2010; Natalia et al., 2012; Tagliazucchi et al., 2012; Liu & Duyn, 2013).
2. **Repeated or quasi periodic spatial temporal patterns (QPP),** which identifies a unique spatiotemporal pattern that is particularly prominent in the default mode network (DMN) and the task positive network (TPN) (Majeed et al., 2009, 2011; Thompson et al., 2014; Belloy et al., 2018; Yousefi et al., 2017).
3. **K-means clustering on windowed functional connectivity,** which identifies discrete periods in time when the spatial patterns of correlated brain activity are relatively stable (Allen et al., 2012)
4. **Recurrence quantification analysis (RQA),** which identifies repeated spatial signatures as a function of time (Webber et al., 2015).

BNMs are likely to perform best at replicating dynamics that are closely tied to the underlying SC of the network and less well for dynamic metrics that are driven by other factors (external input, arousal systems, local connectivity). Thus, the application of dynamic analysis to simulated BNMs can provide insight into which components of the spatial temporal activity can be explained by the structure of the network, and which require a more realistic and complex model. The latter dynamic metrics may lay the foundation for future modeling work that can provide more realistic whole brain simulations.

## Results

### Comparisons to Average Functional Connectivity

We first demonstrate that the models we simulate reproduce common metrics in brain network modeling: average FC and power spectrum. Average functional connectivity matrices were estimated using Pearson correlation pairwise across all ROIs (Figure 1). The ordering of the ROIs, shown in supplementary Table 1, is presented as in Cabral et al., 2011. The Kuramoto and the Fokker-Planck simulations and parameters are very similar to the original model described in Cabral et al., 2011, 2012, but we used a different structural connectivity averaged from the individuals that we had rs-fMRI scans to compare against and it resulted in slightly different parameters (see Methods). To quantify the similarity between the simulated FC matrices and the empirical FC matrix, we calculated the correlation between the two, a method used in previous studies (Cabral et al., 2011). Correlation was 0.37 between Kuramoto and rs-fMRI FC matrices, and 0.5 between Fokker-Planck and rs-fMRI FC matrices. These are in the range of values reported in earlier literature in other BNMs [0.3, 0.7] (Senden et al., 2017; Cabral et al., 2011). Power spectra were calculated for each ROI independently then averaged (Figure 1 bottom right). When plotted on a log-log plot, the BOLD signal has a characteristic 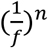 distribution. The power exponent has been reported in literature as 0.88, comparable to the 0.9 measured here for empirical rs-fMRI (Bullmore et al., 2000). The empirical slope falls well within the distribution of the simulated power spectrums. The two simulated models had a slope of 0.74 (Kuramoto) and 0.7 (Fokker-Planck), comparable to a previous report of 0.78 using a different BNMs (Ritter et al., 2013).

**Fig 1.**
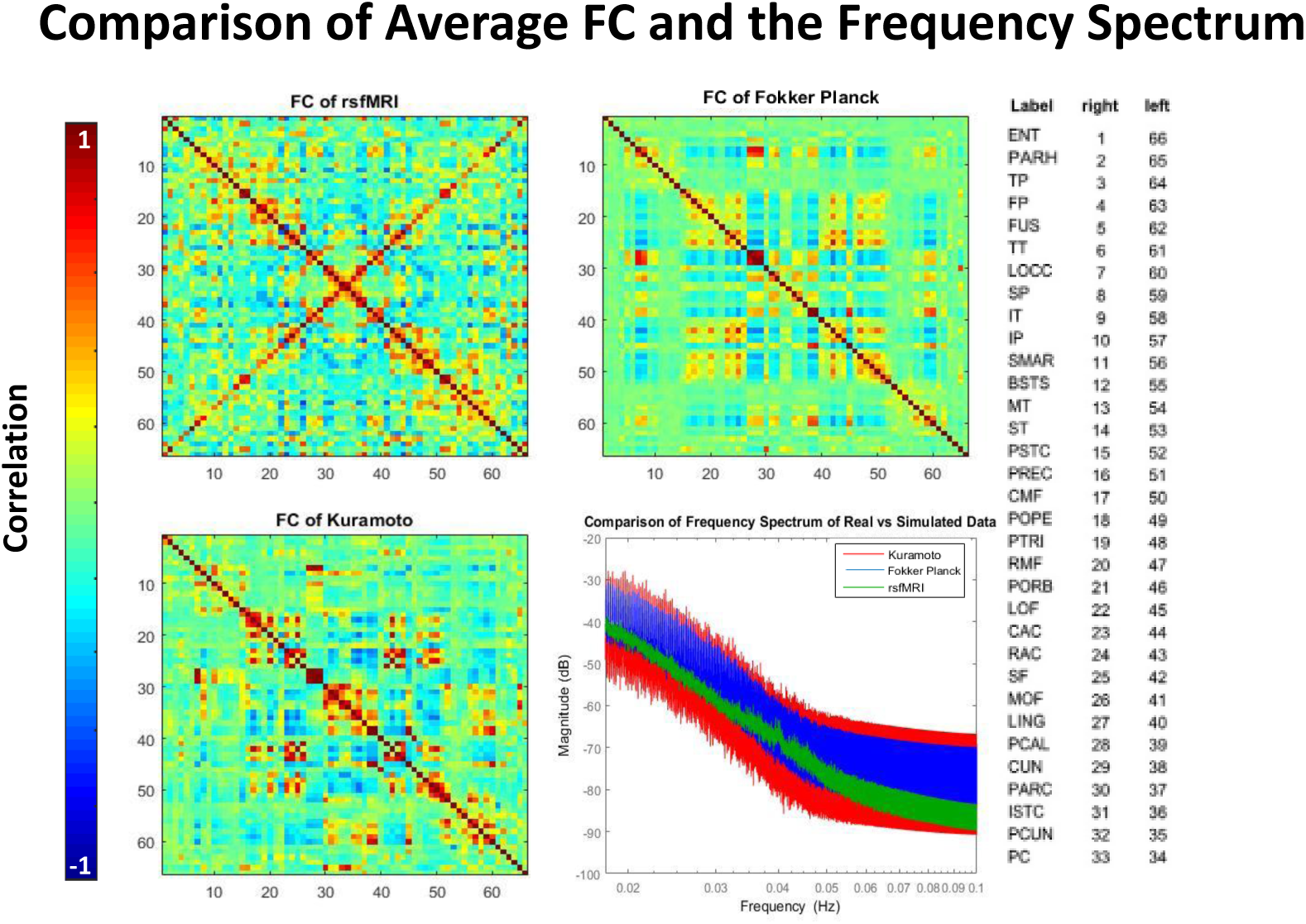
Comparison of the average functional connectivity between the rsfMRI signals and the two simulated models. Correlation between the matrix for empirical data and the Fokker-Planck simulation is 0.5; correlation between empirical and Kuramoto matrices is 0.37. Both modeled matrices and the empirical data exhibit similar structure such as the coordination between hemispheres. The mean frequency spectrum of all ROIs is plotted bottom right and shows that the real signal falls within range of both models. All power spectra exhibit a 1/f^n^ trend.

**Table 1.**

| Parameter name | Differential Equations which were Simulated via Euler Method with a time step of 0.1 ms | State variable describing the neural mass | Cnp - Weight between nodes $n$ and $p$ | Tnp - delay time between nodes $n$ and $p$ | on - SD of gaussian white noise | $\omega_n$ - Oscillator frequency | $c_1$ - First eigenvalue of weight matrix | $\tau_0$ - Relaxation constant |
| --- | --- | --- | --- | --- | --- | --- | --- | --- |
| Fokker Planck | $\tau_0 \frac{dr_n}{dt} = -r_n(t) + \frac{k}{c_1} \sum_{p=1}^N c_{np} r_p(t - \tau_{np}) + \sigma n(t)$ | $r(t)$ - mean firing rate | $k = 0.95$ (Scale of Cnp) | 11 ms set for $\langle Tnp \rangle$ | 2 rad/s | N/A | Calculated | 20 ms |
| Kuramoto | $\frac{d\theta_n}{dt} = \omega_n + k \sum_{p=1}^N c_{np} \sin(\theta_p(t - \tau_{np}) - \theta_n(t)) + \sigma n(t)$ | $\theta(t)$ - oscillator phase | $k = 13$ (Scale of Cnp) | 11 ms set for $\langle Tnp \rangle$ | 2 rad/s | Randomly initialized as $N \sim (60 \text{ Hz}, 2 \text{ Hz})$ | N/A | N/A |

### Point Process/ Neural Avalanche

In coactivation analysis based on the point process approach, all ROIs that cross a certain activation threshold (see methods) (Tagliazucci et al., 2012) are examined at each time point to identify coactivation patterns. Figure 2 shows the coactivation data obtained for the Kuramoto simulation, the Fokker-Planck simulation, and empirical rs-fMRI data. Each value in the matrix represents the fraction of co-occurrences between two ROIs. The matrices are split along the diagonal to show two different threshold values: the top right one at a threshold equaling the mean, and the bottom left at a threshold equal to one standard deviation away from the mean. Both halves of the resulting co-occurrence matrices are highly correlated, greater than 0.9, with their respective functional connectivity matrices calculated via pair wise correlation for both the simulated and empirical modalities (Fig 1). However, at different thresholds the apparent magnitude of coordination between brain regions is different. The cooccurrence rates with a threshold at one standard deviation (bottom half) show a larger difference in connectivity between local networks and global networks than the co-occurrence rates that occur at the mean (top half). The higher values in the top right of the matrix also means that the ROIs are globally more coordinated when they are all cross the mean together. This trend persists at thresholds one standard deviation below the mean (Supplementary Figure 2). Overall, the measure seems to be related to average FC, due to the consistency between the average functional connectivity matrices and the coactivation matrices, but there seems to be a difference on how prominent the local networks vs global coordination plays a role depends on the activity of the signal.

**Fig 2.**
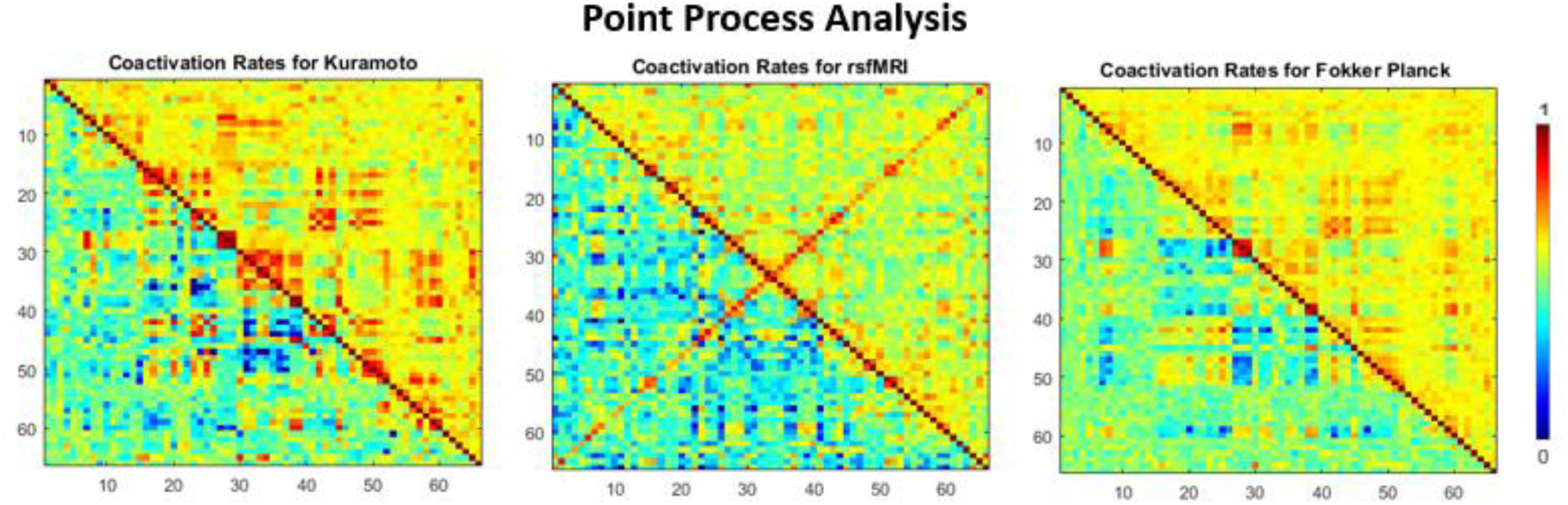
Comparison of the coactivation rates between rs-fMRI, Kuramoto and Fokker-Planck signals. These coactivation rates are calculated by counting the number of co-occurrences of threshold crossings between two ROIs. The log of the rates is plotted. The upper half of the triangle is for the threshold set at the mean, which is zero for the signal. The bottom half of the triangle is for the threshold set at a standard deviation of 1.

### Quasi Periodic Pattern Algorithm Comparison

The quasi periodic pattern (QPP) finding algorithm estimates a recurring spatiotemporal pattern that occurs throughout resting and task states. It consists of a characteristic pattern dominated by the activation and inhibition of the regions that correspond to the DMN and TPN in a specific temporal sequence (Majeed et al., 2009, 2011; Yousefi et al., 2017). The QPP templates obtained from the real data and from each simulation are shown in Figure 3 in a simplified format, where the color bar shows the level of activation or deactivation in each ROI as a function of time. For better visualization, please see the supplementary videos that show the pattern as it evolves over a surface representation of the brain. The pattern in the rs-fMRI data is consistent with the QPP templates obtained previously (Majeed et al., 2011; Yousefi et al., 2017). The QPPs from the two models are very similar to each other (correlation of 0.81), but have important differences from the empirical QPP (correlation of 0.34, 0.33). In fact, the pattern in the simulated models seems to indicate a simple flip between two states, where a subset of ROIs are first active and then inactive. The boxy nature of the plot is due to the spatial ordering of the ROIs that was originally defined by their sub-network connectivity, suggesting that these subcomponents are activating and deactivating together. The QPP obtained from the real data is more complex and demonstrates time lags between areas in addition to changing states. Rather than activating half the network and then deactivating, the empirical QPP suggests that the power in the bold signal cyclically flows through a certain order of ROIs. The relative lengths of the simulated and the observed patterns are different as well. The QPP from the real data is approximately 20 s in length, in agreement with previous reports (Majeed et al., 2011; Yousefi et al., 2017). In contrast, both of the models give QPPs that are ~12-13 s in length, despite the use of identical windows and the similar frequency content of the signals. Histograms showing the correlation between the QPP and the scan for the real, Kuramoto, and Fokker-Planck data indicate that the real QPP exhibits more occurrences of strong correlation or anticorrelation with the scan than either model. Histograms of the correlation for the two models are nearly identical.

**Fig 3.**
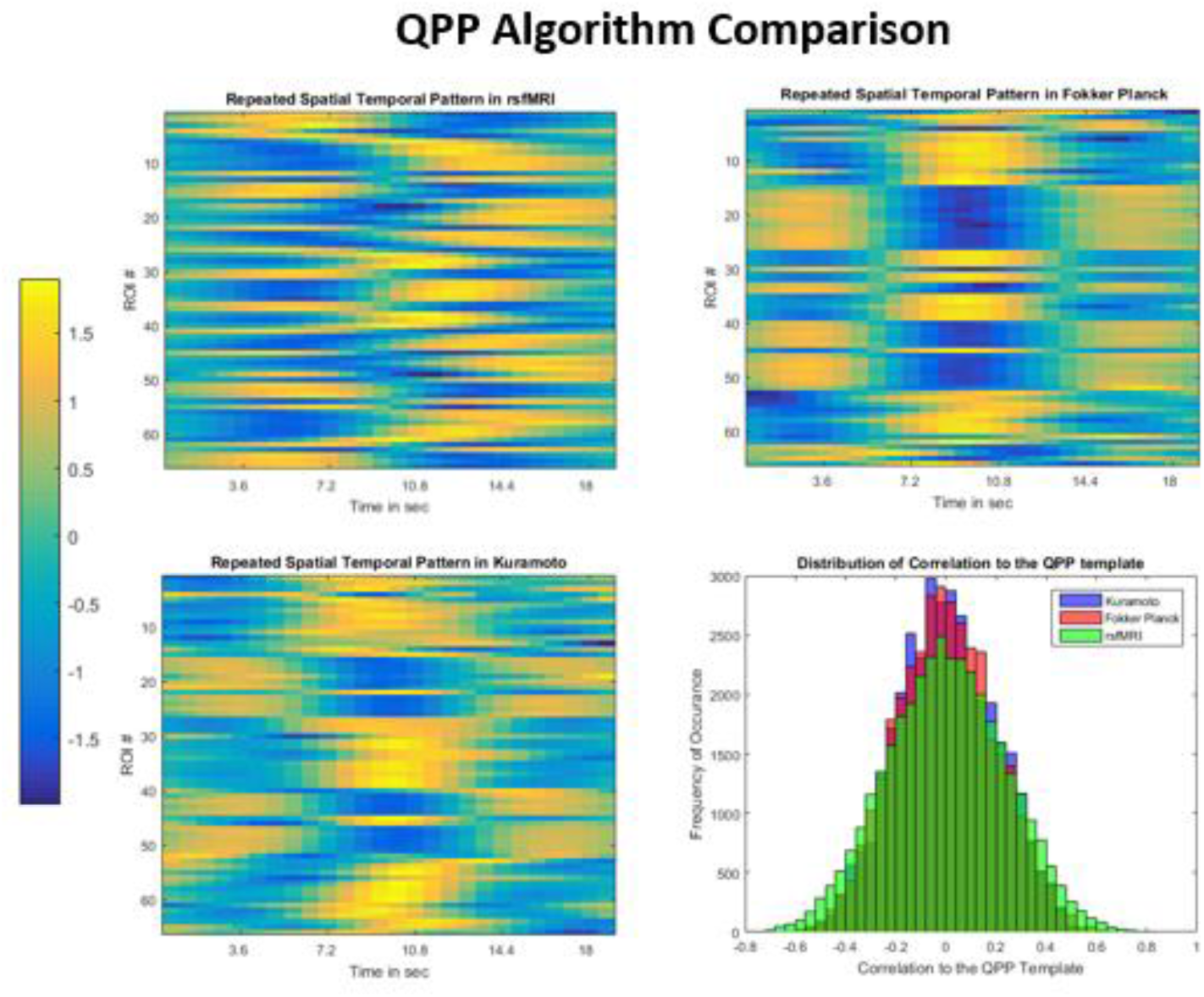
Comparison of the QPPs obtained for each model and the real data. The two simulated models (top right and bottom left) produce templates that are similar to each other but less similar to the template extracted from rs-fMRI. The correlation values to the template are plotted in a histogram (bottom right), shows that the real signal has more extreme values than either model. All three are significantly different (p < 0.0001) in a Komogorov-Smirnov test.

### K-means on Sliding Windowed Matrices

To identify states in the windowed functional connectivity, which is plotted over time in supplementary video 2, we used k-means analysis to compare the real and simulated data. After k-means clustering (k=7), we examined both the spatial composition of the resulting clusters (or states) and metrics that describe how the brain transitions between them (Allen et al., 2012). The corresponding centroids are shown in supplementary Figure 2. The top row in Figure 4, quantifies for each individual scan or simulation, starting at different initial conditions (N= 30), how many of these states are visited, how long they dwell in each state, and how far apart these visited states are on average. The Fokker-Planck and the rs-fMRI have each an average of five states per individual scan, and transition relatively at the same rate, but the average distance is almost twice in the rs-fMRI states compared to the simulated model. Visually the Fokker-Planck centroids look very similar (supplementary figure 3), suggesting that the diversity of states encountered is still very low. The Kuramoto model, produces states whose distance between each other are in the range seen in the real data but each instantiation seems to have fewer states compared to rs-fMRI. Most instantiations (66 percent) result in only a single state, but under certain initial conditions the Kuramoto model can be simulated in order to transition between two to three different states. The model also seems to dwell in these states longer than in the rs-fMRI and Fokker-Planck, where the order of time is also larger than the sliding window length used to calculate the functional connectivity.

**Fig 4.**
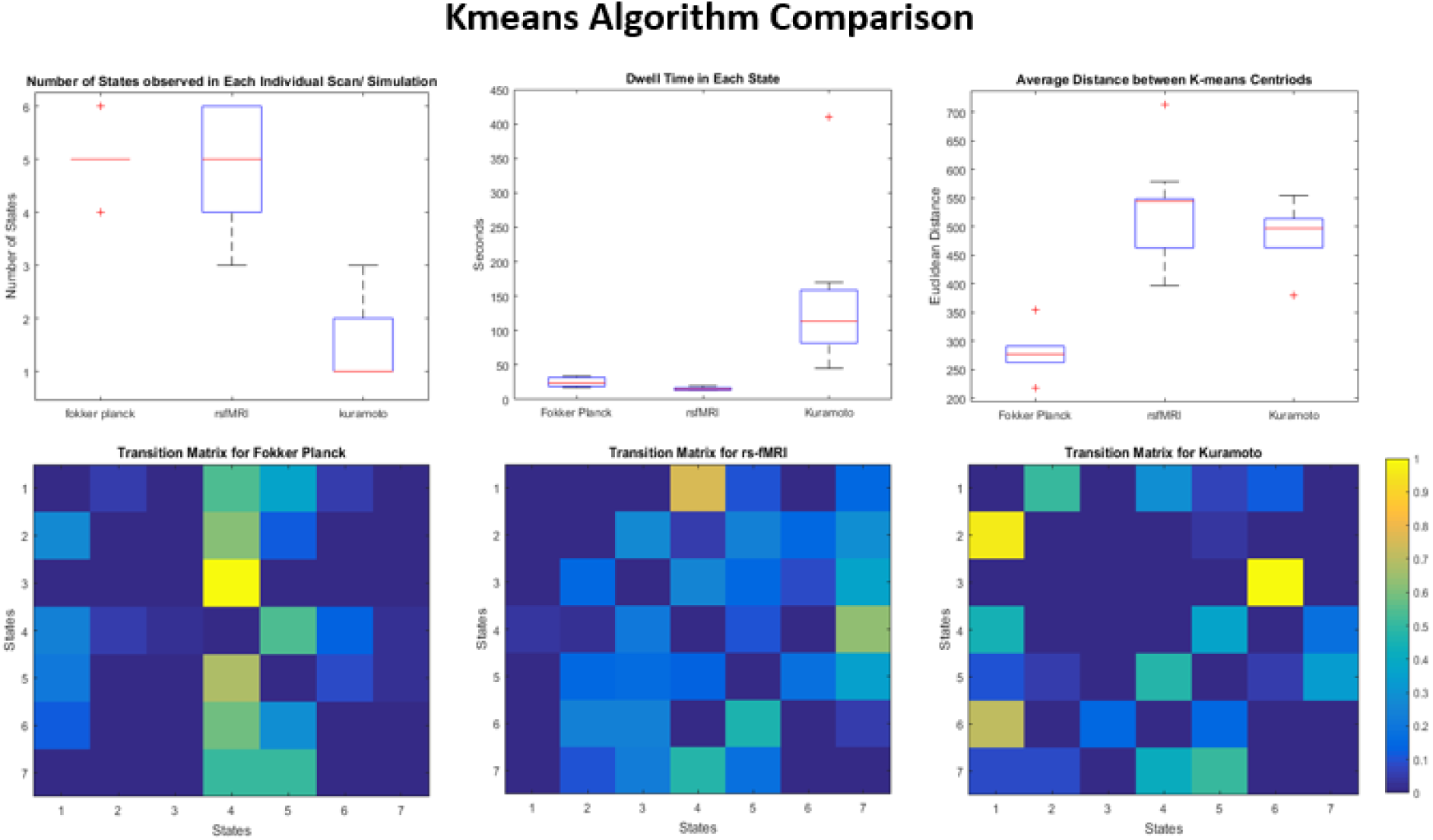
The top row shows the average number of states seen in an individual run (top left), the average dwell time in each state (top middle), and the mean distances between the centroids (top right). The transition matrices between the different kmeans centroids is shown on the bottom row. The values reflect the number of raw occurrences seen in the scan and are not normalized.

The transition matrices for the empirical data also seems to be more complex than in the simulated models (Figure 4 bottom row). The empirical rs-fMRI data has more transitions between states than in either simulated model. The Kuramoto and the Fokker-Planck are roughly around the same complexity (Fokker Planck has 19 off-diagonal zeros, Kuramoto has 20, rs-fMRI has 8), but the Kuramoto model has far fewer states. The Fokker-Planck model thus exhibits the least diverse transitions between states activity. The results are similar to previous studies, where the fast oscillator Kuramoto model has been shown to produce complex state transitions and the Fokker-Planck has been shown to be the naïve model showing no complex dynamics (Hansen et al., 2014, Cabral et al., 2017). In short, neither of the BNMs exhibit the rich set of state transitions as observed with rs-fMRI.

### Recurrence Quantification Analysis (RQA)

Figure 5 shows the reccurance plots for three individual graphs for the empirical and simulated BOLD signal. These plots are calculated by correlating the pattern of activity at each time point with the pattern from every other time point. Diagonal lines that are parallel to the main diagonal represent repeating transistions that are seen throughout the scan, whereas vertical or horizontal blocks represent dwell periods during the scan. A cursory inspection of the three recurrance plots (Figure 5 top row) show that the two models have far less repeating structure than seen in rs-fMRI. This relation is quantified by the bottom three plots that shows the Recurrace Rate (left), Entropy of diagonal lines (middle) and the average length of diagonal lines (right). The reccurance rate seems to be much higher in the Kuramoto and empirical signal than in the Fokker-Planck model. However, the entropy and length of the lines (related to how different the states are and how long they linger) clearly separate the three data sets (bottom and right). Entropy and line length are highest in the real data and lowest in the Fokker-Planck simulation, with the Kuramoto simulation residing in between. The low values for the Fokker-Planck data are likely to be related to the relatively stable configuration of functional connectivity over time as observed with the sliding window correlation analysis. Overall the technique is able to separate the emprical data and the models pretty robustly, and shows a clear difference between the more simpler Fokker-Planck model and the more complex Kuramoto model.

**Fig 5.**
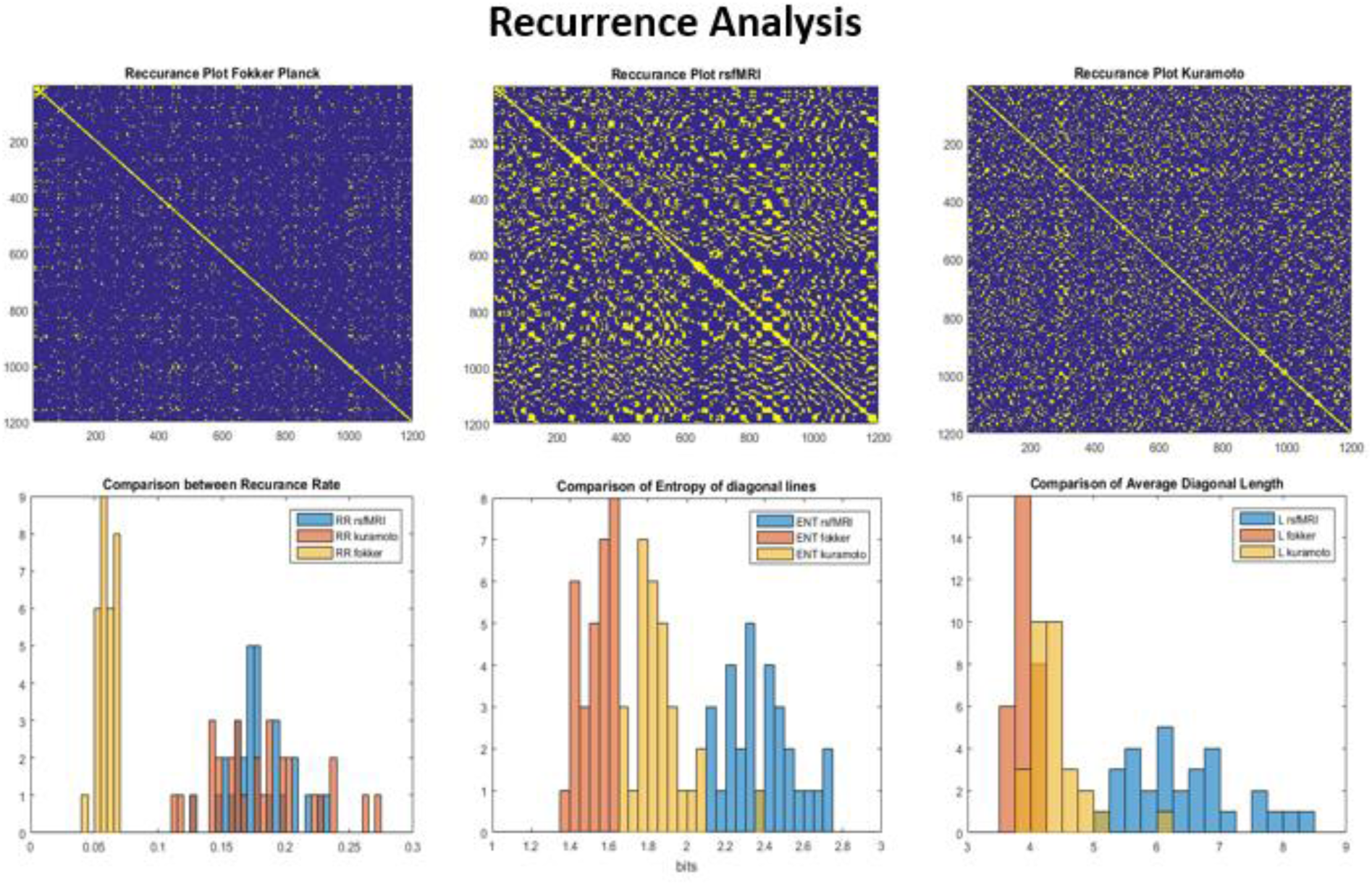
Comparison of the recurrent quantification analysis (RQA) between the simulated and real data sets. The top row shows three scans of recurrent plots thresholded at 0.3 for the three datasets. Bottom row shows the distribution of three different RQA techniques over all scans. Bottom left shows the recurrence rate which is much higher for the Kuramoto and rs-fMRI model than the Fokker-Planck simulation. The recurrence rate is a measure of repeated states seen in the dynamics of the rs-fMRI signal. The middle and the right plots quantify how much similar trajectories occur during the scan. The measured rs-fMRI signal shows much more variance between different trajectories and are of much longer

## Discussion

Our exploratory application of dynamic metrics to simulated BNM data and empirical rs-fMRI data uncovered a number of intriguing similarities and differences, both between the models and real data and between the two models. These findings provide insight into possible mechanisms that underlie empirical network dynamics as well as support the use of dynamic metrics for stronger validation and constraints of BNMs.

### Functional Connectivity and Power Spectra

From previous studies using multiple models and parameterizations, it appears that certain properties of the simulated signal are due to the underlying structural connectivity rather than the model of activity used at each node (Bullmore & Sporns 2009; Stam et al., 2016). In our study, these properties should be similar across models (which share identical structural connectivity) and in the real data. Average functional connectivity analysis is one of these properties. A correlation between the structural connectivity matrices derived from diffusion tensor imaging and the respective functional connectivity estimates from resting state fMRI have a correlation value of 0.45 as measured through our methodology, which is similar to what has been described before (Bullmore & Sporns 2009). In fact, all three dynamical systems produce signals where functional connectivity is highly correlated with the structural input, leading to suggest that average functional connectivity is closely related to the underlying connectome. There are known differences between the two where edges are the result of a third process that drive two structurally disconnected regions, but the majority of the edges between the ROIs can be described as a function of how many white matter tracks run between them (Stam et al, 2016). Moreover, the frequency spectrum and the characteristic 1/f^n^ distribution are similar for both BNMs and the empirical FC, suggesting that the spectrum is a property of the underlying structure of the network.

## Coactivation Patterns

The coactivation analysis also showed a shared feature across the empirical and simulated data. For all three data sets, coactivation patterns were strongly correlated with average functional connectivity. This is not surprising in the real rs-fMRI data, where the low SNR means that small-amplitude changes in the BOLD signal are likely to be indistinguishable from noise. Thus, only relatively high amplitude events are able to consistently emerge from the noise and account for much of the correlation across areas. The similar findings for the BNMs suggest that the simulations accurately capture this property of the BOLD signal.

From the modeling perspective, the role of the network is most pronounced when the signal is at the high level. This can be explained by the original formulation of a the BNMs from Table 1, where the network term would play larger role in the local dynamics of the populations at high levels of its input network activity. This would give arise to a discrete time point where the network activity becomes larger at a certain threshold than the local dynamics. This asymmetric relationship between the network term and the local term is common in the general formulation of BNMs, and could be a property of the unknown empirical dynamical system.

### QPPs

The successful detection of quasiperiodic patterns in both BNMs indicates that these network models capture at least some of the dynamical features of the brain’s activity. The surprising similarity of the QPP templates obtained from the two very different dynamical models suggests that these patterns are a resonance of the structural connectivity of the network. On the other hand, there are substantial differences between the real QPP and the ones obtained from the BNMs. The real QPP is more complex, with gradual switching at different time lags in different areas. This may be a feature of the physical proximity of the ROIs. In QPPs obtained from rs-fMRI data, activity tends to propagate along the cortex as key areas activate or deactivate. The incorporation of aspects of neural field models (Sanz Leon et al., 2015) into the existing BNMs may result is a more accurate reproduction of this propagation. The real QPP template is also longer in length than the ones from the BNMs, despite the similar frequency content of the BNMs and real data, the use of identical preprocessing strategies, and the application of the standard 20 s window during QPP analysis. It is possible that the difference in QPP length results from the unidirectionality of certain white matter connections or other-Properties that cannot be captured using standard tractography.

### Sliding window correlation and state transitions

Some dynamic properties appear to arise more from the complex interactions linked to the unique temporal description of activity in each ROI than from the underlying structural connectivity. These properties are likely to be different for each BNM, and either or neither may be similar to the empirical data. The k-means algorithm on the windowed FC matrices revealed a complex network of states in the rsfMRI data that each demonstrated distinct spatial patterns of connectivity between ROIs and a complex web of transitions between them. In the Fokker-Planck model, there are a similar number of states, but the states are very similar to each other. Seven states were used for all three datasets to be consistent, but it may be more appropriate to consider the Fokker-Planck states as representing a single state artificially divided into multiple components. Our state description in Fokker-Planck, also known as general linear model, echoes previous finding because it is known that we cannot produce stable attractor states with just a linear set of differential equations (Hansen et al., 2014; Cabral et al., 2017). The Kuramoto model for some initial conditions, on the other hand, produced state transitions between states that are as spatially as distinct as the rs-fMRI, but had much fewer states and a simpler transition matrix than the empirical signal. The model tend to dwell in these states for a longer time as well than seen in the real data. This suggests that BNMs can reproduce at least some of the dynamic states observed in rs-fMRI, although current models do not recapitulate the rich variety observed in empirical data.

### Recurrence analysis

In some sense, recurrence analysis captures lingering states similar to the clustering analysis, so it is not surprising that it depends more on the dynamic description of the model than on the underlying structure. Again, the Fokker-Planck model exhibited the least complexity, the rs-fMRI data exhibited the most, and the Kuramoto model fell in between the two. The empirical data has repeated trajectories that occur more often and are longer than each of the simulated BNM. The RQA metrics separate the three models the most effectively as it looks for-Patterns in the time domain, whereas the earlier functional connectivity analysis that examines the spatial domain has the greatest overlap. Average FC and Recurrence analysis both use correlation, except that one uses the space across rows that spans the ROIs whereas the other uses the space of the columns that represent single time points in the BOLD data. Average FC which examines coordination between ROIs reveals a static network related to the input SC. Recurrence analysis that examines the time domain, reveals properties that seem to be most unique to the formulation of each BNM.

### Limitations

Our modeling approach makes many simplifying assumptions that do not capture the true complexity of the brain. In the construction of the structural connectome, we assumed that all connections were bidirectional. This is a limitation of using tractography to build the structural network, since tractography cannot distinguish unidirectional connections. Moreover, estimates of fiber density for connections between regions that have very sharp angles or between regions that are spatially far apart are far lower than the true connectivity between these regions (Bullmore & Sporns 2009). In our generative models we also assumed a homogeneity in the response of ROIs, both in their neural description, as well as their transformation using the hemodynamic Balloon-Windkessel model. Moreover, we did not simulate subcortical structures that are known to play a crucial role in the operation of the central nervous system. All these factors might change the association between dynamic metrics and the simulated BNM signal.

We also examined only a single parameterization for only two BNMs. There are a variety of BNMs, some of which are likely to exhibit more complex dynamics than either the Kuramoto or Fokker-Planck models (Sanz Leon et al., 2015). Even different parameterizations of a single model can give rise to vastly different behavior (Hansen et al., 2014). We chose to focus on the Kuramoto and Fokker-Planck models due to their relative simplicity, their thorough characterization, and the expectation that they would have dissimilar dynamic properties.

There are also numerous dynamic analysis methods available for rs-fMRI (Keilholz et al., 2017). We chose to focus on a few of the most common ones, but future work should certainly examine the use of other types of analysis to produce even more sensitive metrics.

### Conclusion

The analysis of BNMs using dynamic techniques designed for rs-fMRI strengthens the notion that certain properties seem to be a description of the underlying structural connectivity and other-Properties are a function of the specific description of the evolution according to each BNM. The structural properties are most apparent when estimating average functional connectivity or identifying the contribution of individual events, as well as naturally gives arise to repeated spatiotemporal patterns. The temporal properties that distinguished each BNM were most pronounced in metrics that examined different repeated brain states and transitions between them. Using this metrics we can quantify the Kuramoto model performed better than the Fokker-Planck model, although it did not exhibit the level of complexity observed in the real data. The results of this work suggest possible modifications to make BNMs more realistic and establishes dynamic metrics as important tools for distinguishing between different models or-Parameterizations. Moreover, it provides insight into the fundamental sources of the dynamics observed in rs-fMRI in terms of structural connectivity or neural dynamics.

## Methods

### General Methods

The general overview of our methodology is summarized by Figure 6.

**Fig 6.**
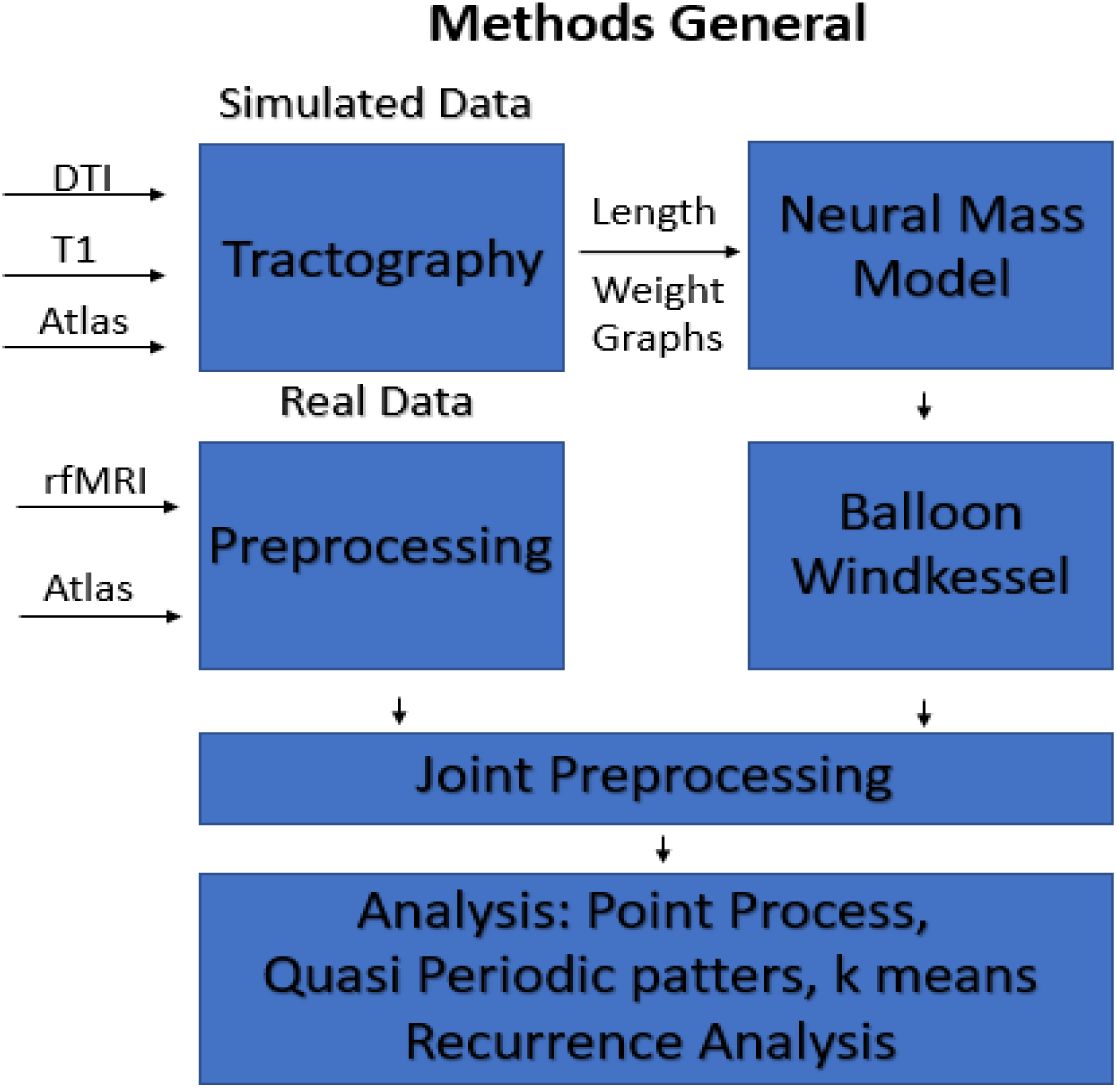
General Workflow - We used DTI data to generate the length and weight matrix between ROIs of our specified Atlas. Using this structural connectome, we generate data using the different Neural Mass Models and transformed into the BOLD signal via the Balloon Windkessel Model. To compare with empirical fMRI data, scans from HCP were preprocessed and parcellated using the same Atlas. The final preprocessing of filtering, global signal regression, and normalization was done jointly for all sets of data. The final data was then analyzed with each of the different dynamical analysis techniques.

### Structural Connectome

Using Human Connectome Project’s Diffusion Weighted Images (Spin Echo TR 5520 ms, TE 89.5 ms, flip angle 78, Voxel 1.25 mm) from 5 random subjects (Van Essen et al., 2013), we generated one average structural connectome. Tractography was performed using the freely available software Mrtrix with maximum fiber length set to 250 mm (J-Donald et al., 2012) and parcellated using the Desikan-Killiany atlas (Desikan et al., 2005). For each subject their respective T1w images (TR 2400 ms, TE 2.14, Voxel Size 0.7 mm) were aligned to the standard space, then the using the warping matrix we transformed the DWI images. Probabilistic Tractography then was run between each ROI and then pruned to generate 10 million fibers. To generate the estimates for the length and weight matrices from the tractography, we used the same methodology as Hagmann et al. (2008). The length between two ROIs was defined as the average fiber length of all fibers that went between them, and the weight was the number of fibers going between two ROIs normalized by the surface area of the receiving ROI. The atlas provides 84 cortical and subcortical ROIs, but we selected the same 66 cortical regions as in (Cabral 2011) for comparison to previous work. The resulting matrices are shown in Figure 7, and there are a few important differences between our tractography, the one from Hagmann et al. (2008), and the ideal tractography. Our current tractography due to longer fiber lengths has more interhemispheric connections than the one presented in Hagmann et al. (2008). However, tractography is less sensitive to longer connections (Fornito et al., 2013) and therefore the between hemispheric connections were scaled by a factor of 4 to offset the known issue. Tractography is also less sensitive to fibers with sharp angles than to fibers with more straight angles, so for example it results in less connections between the two primary visual areas (ROIs 27 -29) that have the sharpest bend in the corpus callosum (Fornito et al., 2013). The final weight matrix was normalized using the matrix norm function to be unit norm. The length matrix was divided by the mean conduction velocity 5.45 m/s to get the delay matrix. This set the mean delay to 11 ms in accordance to the Cabral et al. (2011).

**Fig 7.**
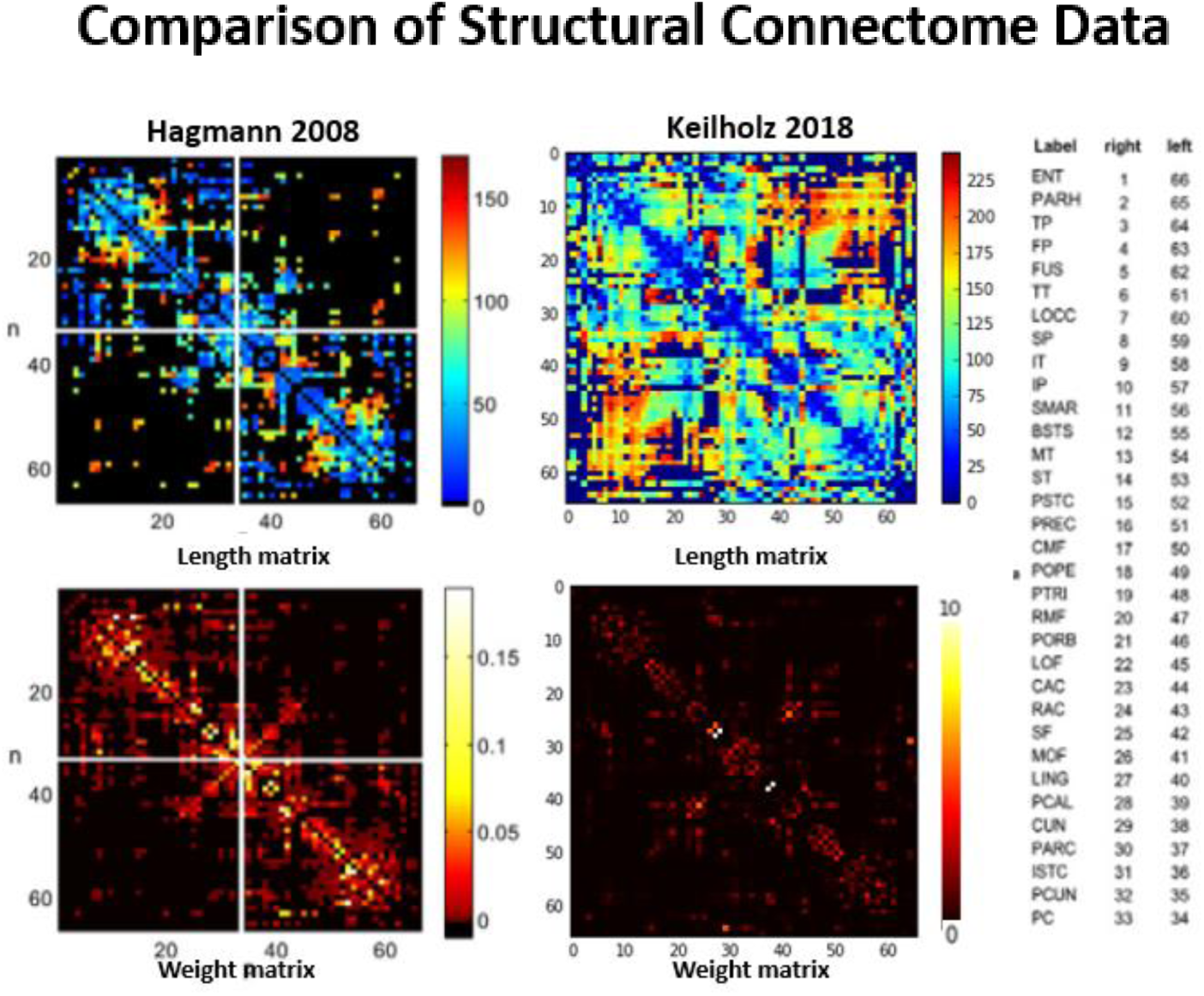
Comparison of our tractography with the tractography that is commonly used by Hagmann et al., 2008. Our tractography has set the max tract length to 250 mm, which allows us to image the longer tracts that are between hemispheres. Top row matrix of the mean length of fiber between two ROI regions measured in mm. Bottom row – number of fibers between two ROI regions divided by the surface area of the receiving ROI (row -> col) and then normalized to one. Left – Hagmans matrices, and right the corresponding matrices from our tractography.

#### BNMs

Brain Network models describe the BOLD signal as the coupling of n distinct neural populations corresponding to different cortical regions. Each population is connected via a weight matrix obtained from structural connectivity that describes the strength of the connection between nodes. In general each of these n areas are modeled by a differential equation for each node: 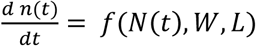 where N(t) is the time series of all the nodes/ROIs, W is the weight matrix, L is the length matrix, and for given random initial conditions for *n*_0_, the timeseries n(t) can be solved for by using the Euler integration method (Sanz et al., Leon 2015). The timeseries n(t) is the state variable and is representative of a measurable property of the neural mass such as firing rate. Some variants use more than one variable to represent the state of the neural mass, but in this paper, we consider two models that only use one state variable, namely the Fokker-Planck and the Kuramoto models. Table 1 shows the mathematical description as well as the values of the parameters used in the simulations.

The Kuramoto model is derived from an assumption that each neural population is in a closed periodic trajectory in phase space that represents its computational processing (Cabral et al., 2011). It has been shown that it can then be modeled by a phasic oscillator can be described by a single parameter theta which represents its location within a 2pi cycle. Inputs into these phasic oscillators perturb its trajectory but it stays within its limit cycle. Each of these oscillators couple via the network and are driven to the same angle and thus synchronizing the oscillators by a function of the difference between the angles of the phasic oscillator.

The Fokker-Planck model assumes that the mean firing rate of the neural populations are distributed in a gaussian manner. They make this assertion in accordance to the Central Limit Theorem, which states that the sum of uncorrelated random processes converges to a Gaussian probability distribution, even if the individual processes are highly non-Gaussian. Inputs into this neural mass shift the mean firing rate to a higher firing rate. The mass shifted from its equilibrium tries to relax at the rate proportional to its own firing rate, keeping the system stable via negative feedback.

For each model, the differential equations were numerically integrated with a time step function of 0.1 ms for a duration of 15 min to match the length of a HCP rs-fMRI scan. The first 20 sec are thrown away to avoid transient effects from initial conditions. The choice for the values for all the parameters given in table 1 follow previous work by Cabral 2012 and Cabral 2014, except that the values for k are slightly different than the ones in the paper to account for differences in the structural connectivity matrix. The values were slightly smaller for the Kuramoto (13 instead of 18) because there were more numerous connections in the newer tractography. Simulations of functional connectivity and the intermediate steps with the original Hagmann’s matrices and the comparisons with Cabral 2011 are given in the supplementary fig 2.

#### Converting to BOLD

In order to compare the neural simulated data with the hemodynamic response measured from fMRI, we have to convert the high frequency activity down to the low frequency hemodynamic response. In order to do this we utilize the Balloon-Windkessel model, which is a quadruple differential equation model that takes in an neuronal input and calculates the blood flow and blood volume and uses that to estimate the fraction of the oxygenated blood to the deoxygenated blood (Stephan et al., 2007; Friston et al., 2003). Supplementary Figure 3 shows the impulse response of our Balloon-Windkessel model which looks roughly like the canonical hemodynamic response function. We used the same constants for our Balloon Model as those given in Friston et al. (2003). After-Passing the output of the BNMs through the Balloon-Windkessel model, we then downsampled to the same sampling rate as the rs-fMRI data (0.72 s).

#### Pre-processing rsfMRI

For the rsfMRI data we used 30 individual HCP scan (Gradient echo EPI, TR 720 ms, TE 33.1ms, flip angle 52, Voxel 2mm) that are each roughly 15 min long. The data came from the minimally processed pipeline and then was ICA denoised using the 300 ICA vectors that HCP provides. We then applied the same Desikan-Killiany atlas as used in the tractography onto the data and obtained the mean time series for each ROI. From then on, we applied the same processing pipeline for the simulated data as the real data, in order to keep the processing as similar as possible. These steps in order were z-scoring each time series, then band passing filtering the signal from 0.01 to 0.25 Hz, and then global signal regression using a linear regression model, and then applying a final z-score step. These steps were selected in accordance with Cabral et al. (2011).

#### Dynamic Analysis Techniques

To compare the dynamics of the rs-fMRI signal and the BNMs, we selected analysis techniques that are commonly used and characterized the signal at different spatial and temporal scales. Table 2 shows a quick comparison of the different techniques that were applied.

**Table 2.**
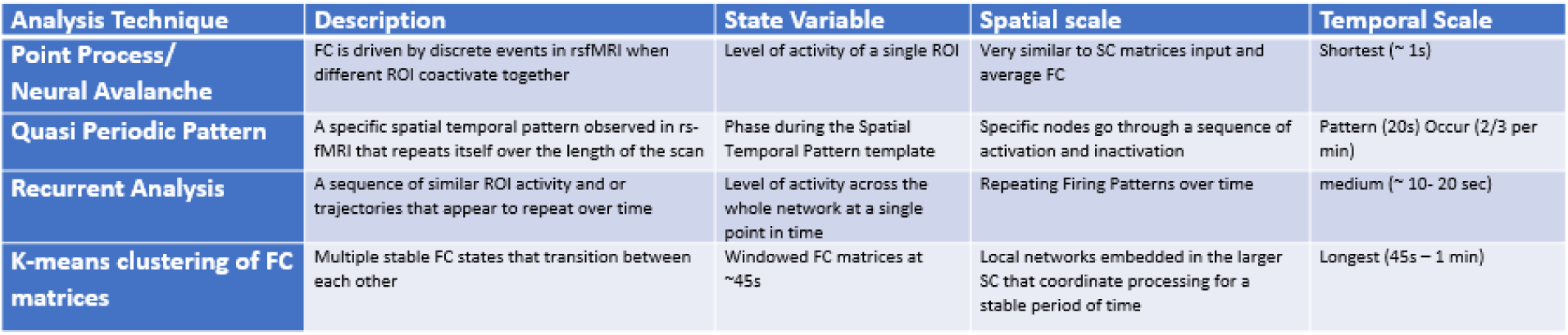

The Point Process assumes that activity in an area triggers neural avalanches in regions that are involved in the information processing (Tagliazucci et al., 2012). The signal is only interpretable in either at its high levels of activation or very low levels of activation when it is coordinating information transfer with other elements in the network. Later models explicitly write out the mathematical formulation using impulse response and solve for a sparse representation of these coactivation patterns which are thought to be unique computational trajectories across the brain (Karahanoğlu et al., 2015; Liu & Duyn 2013). But in this analysis, we use Tagliazucci’s methodology by quantifying when different ROIs cross the same threshold over time. We implemented this approach by recording when the activity at a certain ROI crosses a certain threshold and then counting how many other ROIs cross the same threshold within 3 timesteps (0.72s) of the original crossing. We normalize the co-occurrence rates to get a fraction by dividing by the total number of crossings at each ROI. We applied this analysis with two different thresholds, one at the mean of the signal and one at one standard deviation away which for our normalized signals were at zero and one respectively. Prior work that has shown that average functional connectivity is primarily driven by coactivation events (Tagliazucci et al., 2012).

A second approach examines the quasiperiodic patterns of BOLD signal propagation over the course of the scan. The QPP algorithm identifies the most prominent repeating spatiotemporal pattern in the signal (Majeed et al., 2009, 2011). In brief, the algorithm chooses a random chunk of the rs-fMRI data (20s of data) and correlates it with the entire scan (Majeed et al., 2011). Time points with high correlation to the random chunk indicate repeated occurrences and are averaged together to form the new template. This process is iterated until the template converges. Since we use a random seed point as the original template, repeated runs of the algorithm produce QPPs with different phases. Therefore, in order to compare the patterns from the rs-fMRI and the simulated models, the QPP was circularly shifted to the point where maximum correlation occurred.

Sliding window correlation followed by k means clustering was applied to examine the brain states and transitions in each set of data (Allen et al., 2012). Using a sliding window length of 60*0.72 sec, Pearson correlation was calculated pairwise for all ROIs. The window was then advanced by one time point and the process was repeated until the window reached the end of the scan. This value is around the range used in previous work (Allen et al., 2012). Correlation values were Fisher-transformed to better approximate a normal distribution and the k-means algorithm was applied to cluster the data into 7 groups using Manhattan distance based on previous studies (Allen et al., 2012). Clustering was repeated ten times and the best resulting clustering was chosen based on minimizing the total distance from the cluster centroids and the feature vectors in order to mitigate the effects of randomly choosing the centroid locations.

Recurrent analysis was performed by calculating correlation of the spatial pattern of activity pairwise across all time points. We then thresholded the values at 0.3, based on literature search, and created recurrent plots (Cabral et al., 2014; Basset et al., 2012). These metrics were calculated using freely available Matlab toolbox (Ouyang et al., 2014) and their distribution for each type of data was plotted. Recurrence Rate, Entropy Rate, and average Diagonal Length were measured. The Recurrence Rate is the rate that similar states occur throughout the scan. Entropy Rate quantifies the difference between repeated states. The average Diagonal Length measures how long these trajectories occur. Collectively they give us an insight of how often similar states occur, how different they are from each other, and how long each of these spatial-temporal trajectories persists.

